# Standard rodent diets differentially impact alcohol consumption and preference and gut microbiome diversity

**DOI:** 10.1101/2024.02.06.579237

**Authors:** Aline Zaparte, Evan Dore, Selby White, Franciely Paliarin, Cameron Gabriel, Katherine Copenhaver, Samhita Basavanhalli, Emily Garcia, Rishith Vaddavalli, Meng Luo, Christopher M. Taylor, David Welsh, Rajani Maiya

## Abstract

Alcohol Use Disorder (AUD) is a complex and widespread disease with limited pharmacotherapies. Preclinical animal models of AUD use a variety of voluntary alcohol consumption procedures to recapitulate different phases of AUD including binge alcohol consumption and dependence. However, voluntary alcohol consumption in mice is widely variable rendering it difficult to reproduce results across labs. Accumulating evidence indicates that different brands of commercially available rodent chow can profoundly influence alcohol intake. In this study, we investigated the effects of three commercially available and widely used rodent diet formulations on alcohol consumption and preference in C57BL/6J mice using the 24h intermittent access procedure. The three brands of chow tested were LabDiet 5001 (LD 5001), LabDiet 5053 (LD 5053), and Teklad 2019S (TL2019S) from two companies (Research Diets and Envigo respectively). Mice fed LD5001 displayed the highest levels of alcohol consumption and preference followed by LD5053 and TL2019S. We also found that alcohol consumption and preference could be rapidly switched by changing the diet 48h prior to alcohol administration. Sucrose, saccharin, and quinine preference were not altered suggesting that the diets did not alter taste perception. We also found that mice fed LD5001 displayed increased quinine-resistant alcohol intake compared to mice fed TL2019S, suggesting that diets could influence the development of “compulsive” like alcohol consumption. We profiled the gut microbiome of water and alcohol drinking mice that were maintained on different diets and found significant differences in bacterial alpha and beta diversity, which could impact gut-brain axis signaling and alcohol consumption.

## Introduction

Alcohol use disorder (AUD) is a persistent and recurring brain disorder marked by a diminished capacity to cease or regulate alcohol consumption, even in the face of negative repercussions (Koob and Volkow, 2016). Alcohol is one of the most misused drugs worldwide, and within the United States, AUD has a lifetime prevalence rate of nearly 8.6% (Glantz et al., 2020). Alcohol consumption significantly impacts both the central and peripheral nervous systems, and its toxic effects can promote cell and tissue injuries throughout the body (Dubinkina et al., 2017). Currently, there are only three FDA-approved treatments for AUD that all suffer from variable efficacy (Lohoff, 2022). Hence, there is an urgent need for the discovery of novel therapeutics to treat AUD.

Animal models of AUD enable us to identify neuroadaptations that underlie the development of alcohol dependence, thereby enabling the discovery of novel therapeutic targets to mitigate AUD. Preclinical studies rely on a suite of voluntary alcohol consumption procedures to capture different phases of AUD, such as binge alcohol consumption, alcohol dependence, and relapse and reinstatement to alcohol seeking (Rhodes et al., 2005; Tabakoff and Hoffman, 2000; Vengeliene et al., 2014). However, the amount of alcohol consumed in voluntary alcohol consumption procedures often vary widely between laboratories, rendering it difficult to reproduce key findings. A variety of factors could contribute to this variability including housing conditions such as type of bedding, temperature, and humidity in the vivarium (Crabbe and Wahlsten, 2003). However, accumulating literature suggests (Maphis et al., 2022; Marshall et al., 2015; Quadir et al., 2020) that the type of rodent chow could be a significant contributor to variations in alcohol consumption. Rodent chow is not standardized across laboratories and can vary significantly in composition and texture. Diet can profoundly influence behavioral outcomes through a variety of pathways, including signaling through the gut-brain axis (Leclercq et al., 2020) and altering taste perception (Tordoff et al., 2002). Previous studies have examined the effects various commercial rodent diets on alcohol consumption and preference (Maphis et al., 2022; Marshall et al., 2015; Quadir et al., 2020). These studies have found that the type of rodent chow used can markedly affect not only the amount but also the pattern of alcohol intake in voluntary drinking procedures. However, the mechanism(s) by which diet can influence alcohol consumption has not been examined in these studies.

One potential mechanism by which rodent diet could influence alcohol consumption is via modifications to the gut microbial composition. Diet is the main modulator of gut microbiome composition, and consequently can alter gut-brain axis signaling (Lynch and Pedersen, 2016), which may play a role in accelerating the addiction cycle (Bravo et al., 2011; García-Cabrerizo et al., 2021). Alcohol consumption by itself can also alter gut microbial communities and lead to dysbiosis, gut leak, and trigger end-organ chronic inflammation (Stärkel et al., 2018). Additionally, the modifications of the gut microbiota due to alcohol consumption plays a role in the development of alcohol-associated diseases, such as nonalcoholic fatty liver, cardiovascular diseases and neuro-psychiatric disorders (Chang and Kao, 2019). Whereas diet plays a pivotal role in shaping the gut microbiome, (Lynch and Pedersen, 2016) little is known about how food composition and nutritional habits regulate the motivation to seek drugs. Given the influence of the diet on the gut microbiome and the connection between the gut microbiota and behavior, we aimed to investigate the impact of commercially available rodent diets on alcohol consumption and preference, as well as gut microbiome diversity.

## Materials and Methods

### Animals

7-8 weeks old male and female C57BL/6J mice were purchased from The Jackson Laboratory (Bar Harbor, ME). Mice were provided access to food and water *ad libitum* and individually housed. Mice were allowed to habituate to a reverse light/dark cycle (on at 10:00 A.M., off 10:00 P.M.) for 1-week prior to initiation of experiment. During the habituation period, mice were individually housed in double grommet cages equipped with two bottles provided with sipper tubes containing water. Animals were weighed once per week.

### Rodent Diets

Animals were provided *ad libitum* access to specified diets of either Labdiet 5001, Labdiet 5053, (Research Diets, New Brunswick, NJ) or Teklad 2019S (Envigo, Indianapolis, IN). The diets did not differ significantly in terms of the amount of carbohydrate, fat, and protein content. However, there were numerous differences in micronutrient constituents, some of which are listed in Table 1.

**Table 1:** Major similarities and differences in macro and micronutrient composition of the three standard rodent diets used in the study.

| Nutrients | Diet |  |  |
| --- | --- | --- | --- |
|  | TL2019S | LD5053 | LD5001 |
| *Energy* |  |  |  |
| Calories from Carbohydrates (%) | 55.00 | 62.38 | 58.00 |
| Calories from Fat (%) | 22.00 | 13.12 | 13.50 |
| Calories from Protein (%) | 23.00 | 24.50 | 28.51 |
| Metabolizable Energy (kcal/gm) | 3.30 | 3.02 | 3.02 |
| *Fatty Acids* |  |  |  |
| C18:2ω6 Linoleic (%) | 3.90 | 2.32 | 1.22 |
| Fat (acid hydrolysis) (%) | 9.00 | 6.30 | 5.70 |
| Fat (ether extract) (%) | 10.00 | 5.00 | 5.00 |
| Fiber (crude) (%) | 2.60 | 4.40 | 5.10 |
| Total Monounsaturated (%) | 1.70 | 1.00 | 1.60 |
| Total Saturated (%) | 1.20 | 0.77 | 1.56 |
| *Minerals* |  |  |  |
| Copper (mg/kg) | 15.00 | 13.00 | 0.13 |
| Iodine (mg/kg) | 6.00 | 0.97 | 1.00 |
| Manganese (mg/kg) | 80.00 | 82.00 | 0.70 |
| Potassium (%) | 0.40 | 1.07 | 1.18 |
| *Vitamins* |  |  |  |
| Biotin (ppm) | 0.90 | 0.30 | 0.30 |
| Choline Chloride (ppm) | 1200.00 | 1575.00 | 2250.00 |
| Folic Acid (ppm) | 9.00 | 3.00 | 7.10 |
| Niacin (ppm) | 120.00 | 84.00 | 120.00 |
| Pantothenic Acid (ppm) | 140.00 | 17.00 | 24.00 |
| Pyridoxine (ppm) | 26.00 | 9.60 | 6.00 |
| Riboflavin (ppm) | 27.00 | 8.10 | 4.50 |
| Thiamin Hydrochloride (ppm) | 117.00 | 16.00 | 16.00 |
| Vitamin A (IU/gm) | 30.00 | 15.00 | 15.00 |
| Vitamin B12 (mcg/kg) | 150.00 | 51.00 | 50.00 |
| Vitamin D3 (IU/gm) | 2.00 | 2.30 | 4.50 |
| Vitamin E (IU/kg) | 135.00 | 99.00 | 42.00 |
| Vitamin K ( as menadione) (ppm) | 100.00 | 3.30 | 1.30 |

### Two bottle choice 24h intermittent access alcohol (IA) consumption procedure

Mice were subjected to IA alcohol consumption procedure as previously described (Hwa et al., 2011). Animals were given two bottles from which to drink, one containing water and the other containing15% ethanol (V/V) (PHARMCO-AAPER, Brookfield, CT) solution for 24 hours every other day. On the off-days, mice had access to two bottles of water. Alcohol and water bottles were weighed prior to placing in the cage and were weighed again 24h later. Spillage was controlled for by measuring volume lost from alcohol and water bottles placed on an empty cage. The position of the water and alcohol bottles were switched for every drinking session. Control water drinking mice were provided with two bottles of water. Mice were weighed once per week and their body weights recorded.

#### Blood ethanol concentration (BEC) measurement

We measured BECs using Analox AM1 analyzer (Analox instruments, Lunenberg, MA) as described (Avegno and Gilpin, 2019). Briefly, blood was collected from tail veins in heparinized capillaries 3h after alcohol bottles were introduced. Blood was spun down at 9000 X g for 12 minutes and plasma was collected. 5ml of plasma was used for BEC measurement. Single point calibration was done for each set of samples with reagents provided by Analox Instruments.

### Continuous access sucrose, saccharin, and quinine consumption

Mice were given access to sucrose saccharin, and quinine in a two-bottle continuous access procedure as described (Maiya et al., 2021). Mice were first given access to sucrose (4%), followed by saccharin (0.03% and 0.06%) and then quinine (100μM, 175μM, and 250μM). Mice had access to each concentration of sucrose/saccharin/quinine for a total of 48h. Positions of the tastant and water bottles were switched every 24h. There was a one-week interval between the different tastants when mice received two bottles of water. Bottles were weighed prior to placing in the cage and again after 24h to determine the amount of fluid consumed.

### Quinine adulteration of alcohol

Alcohol was adulterated with increasing concentrations of quinine ranging from 100mM to 500mM in the IA procedure. Mice were exposed to each concentration of quinine adulterated alcohol for 48h.

### Sample collection

Stool pellets were collected at two timepoints: 1) immediately before the last IA session before diet switch when mice were on TL2019S diet and 2) and immediately before the 4^th^ session post diet switch when mice were on LD5001 or LD5053 diet. Stool pellets were collected in autoclaved microtubes with attached caps and frozen. Stool samples were sent to the Louisiana State University School of Medicine Microbial Genomics Resource Group, for bacterial quantification and analysis.

### DNA extraction, PCR amplification, and sequencing

DNA extraction and sequencing were performed by the Louisiana State University School of Medicine Microbial Genomics Resource Group (http://metagenomics.lsuhsc.edu/). The genomic DNA was extracted using the QIAamp DNA Stool Mini Kit (Qiagen, Valencia, CA) modified to include bead-beating and RNase A treatment. A negative control was set for checking any potential bacterial DNA existing in chemicals or involving during the DNA extraction process. Two steps of amplification were performed to prepare sequencing library using the AccuPrime Taq high fidelity DNA polymerase system (Invitrogen, Carlsbad, CA). A negative control with the control from DNA extraction and a positive control of ZymoBIOMICS™ Microbial Community Standard (ZYMO Research, Irvine, CA) were set during amplicon library preparation. 16S ribosomal DNA hypervariable region V4 was amplified using 20ng genomic DNA and the gene-specific primers with Illumina adaptors: Forward: 5’TCGTCGGCAGCGTCAGATGTGTATAAGAGACAGGTGCCAGCMGCCGCGGTAA3’; Reverse 5’ GTCTCGTGGGCTCGGAGATGTGTATAAGAGACAG GGACTACHVGGGTWTCTAAT 3’. The PCR conditions were as follows: 95 °C for 3 min, 25 cycles of 95 °C for 30 s, 55 °C for 30 s and 72 °C for 30 s, 72 °C for 5 min and holding at 4 °C. PCR products were purified using AMPure XP beads which the beads was added as 0.85x PCR volume. 4 µl purified amplicon DNA from the last step was amplified 8 cycles with the same PCR condition using the primers with different molecular barcodes: forward 5’ AATGATACGGCGACCACCGAGATCTACAC [i5] TCGTCGGCAGCGTC 3’; reverse 5’ CAAGCAGAAGACGGCATACGAGAT [i7] GTCTCGTGGGCTCGG 3’. The indexed amplicon libraries purified using AMPure XP beads and quantified using Quant-iT PicoGreen (Invitrogen) were normalized and pooled. The pooled library was quantified using KAPA Library Quantification Kit (Kapa Biosystems, Cape Town, South Africa) diluted and denatured as the guideline of Illumina’s sequencing library preparation. 10% PhiX was added to the sequencing library as an internal control and increase diversity of 16S RNA amplicon library. The paired-end sequencing was performed on an Illumina MiSeq (Illumina, San Diego, CA) using the 2 × 250bp V2 sequencing kit. The sequencing reads were transferred to Illumina’s BaseSpace for quality analysis, and the generated raw FASTQ files were used for further bioinformatics analysis.

### Sequence Analysis

Sequenced reads underwent analysis utilizing R (v4.3.1) and DADA2 (v1.26.0) (Refs). Initial preprocessing involved the removal of 20 base pairs from both the beginning and end of each read to eliminate low-quality regions flanking the reads. The DADA2 algorithm was then employed to identify sequence variants, with further trimming of the 5’ ends based on these variants. To enhance the detection of rare sequence variants, reads from all samples were aggregated. Taxonomic classification of sequence variants was accomplished using the Silva database (v138.1). Decontamination procedures were carried out using the decontam package (v1.20) with the prevalence method. Additionally, an abundance filter was applied, excluding sequence variants with a relative abundance below 0.01%.

Bacterial richness was estimated using three indexes (1) Observed features, represent the number of the different OTUs found in the sample (richness); (2) Shannon Index considers the diversity of subspecies (richness) and the relative abundance (evenness) of each subspecies within a specific zone; and (3) Simpson considers both the evenness and the percentage representation of each subspecies within a biodiversity sample in a specific zone. This index operates under the assumption that the relative proportion of individuals in an area reflects their significance to overall diversity (Nagendra, 2002).

While alpha-diversity measures the microbiome composition, the beta-diversity measures the distances between the bacterial communities; the dissimilarities between them. Beta-diversity was estimated using Aitchison distance, and the centered log ratio (CLR) transformed abundances, were used to perform principal component (PC) analysis. The Analysis of the dissimilarity (ADONIS) was performed using permutational multivariate analysis of variance (PERMANOVA) technique and Pairwise ADONIS was applied as a *post hoc* test (R vegan package) (By, 2022)

## Results

We first compared alcohol consumption and preference in male C57BL/6J mice that were maintained on LD5053 or TL2019S diets. Mice maintained on LD5053 consumed significantly more alcohol than their TL2019S-fed counterparts (**Fig. 1A**). Two-Way RM ANOVA of alcohol consumption revealed a significant diet X session interaction [(F _diet X dession_ (7, 133) = 3.722, P = 0.0010)], and a marginally significant main effect of diet [(F _diet_ (1, 19) = 4.268, P =0.052)]. Sidak post-test revealed that mice on LD5053 consumed significantly higher amounts of alcohol during the last drinking session (**Fig. 1A**). Two-way RM ANOVA of alcohol preference indicated a significant diet X session interaction (**Fig. 1B**) [(F _diet X session_ (7, 133) = 6.120, P = 0.0061)]. However, Sidak post-test did not reveal any significant differences between the two groups across individual drinking sessions.

**Figure 1:**
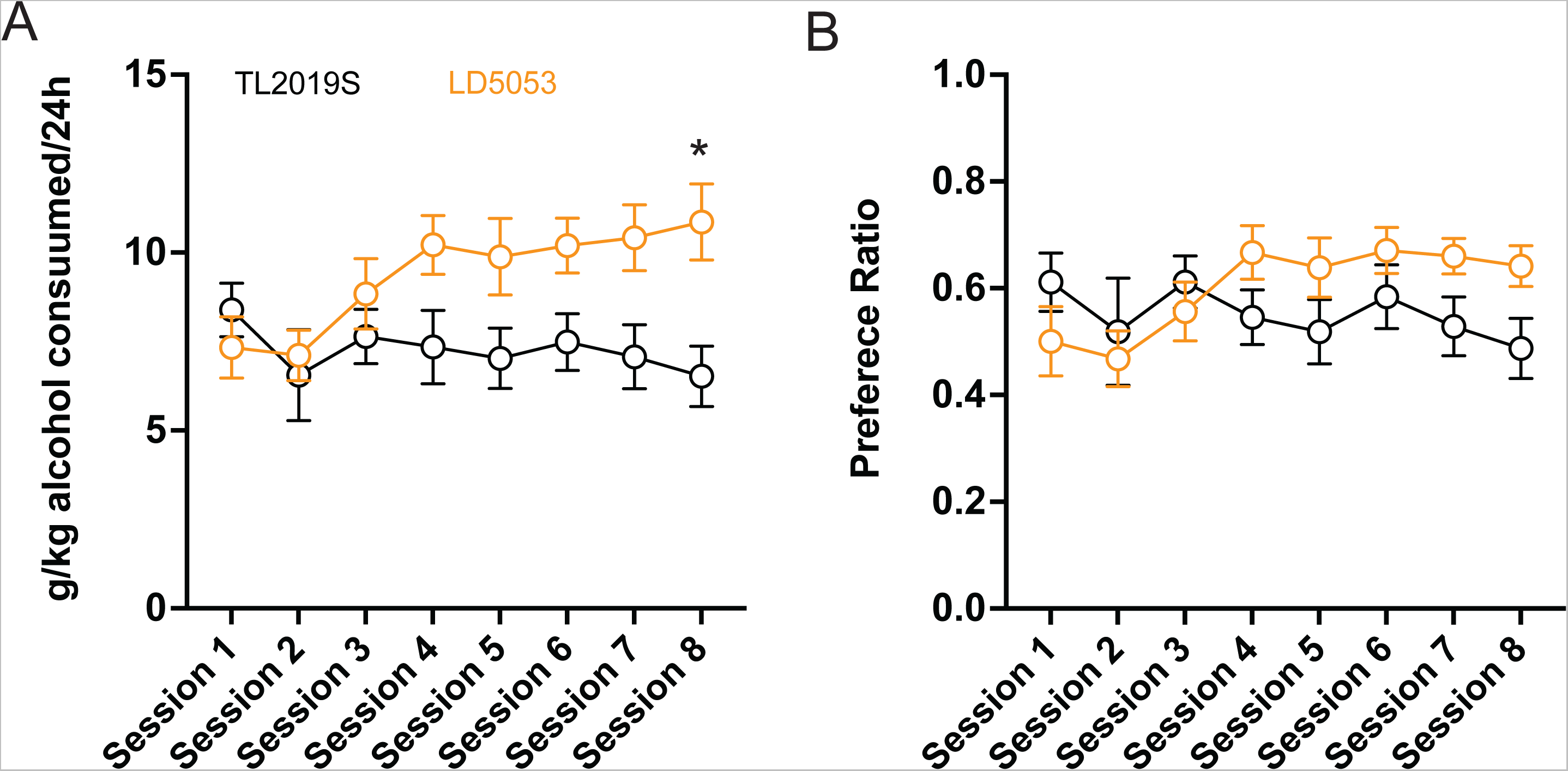
Mice fed LD5053 consumed significantly more alcohol (A) and showed increased preference (B) for alcohol in the IA two-bottle choice procedure. *, P<0.05, Sidak’s post-test, N = 7/group.

We next determined the consequence of switching the diet from TL2019S to LD5053 or LD5001, another commonly used diet formulation from Research Diets, on alcohol consumption and preference. Diets were switched 48h prior to the alcohol drinking session. We first measured the consequences of switching from TL2019S to LD5001 diet. We found a significant increase in alcohol consumption (**Fig. 2A**) and preference (**Fig. 2B**) when mice were switched from TL2019S to LD5001. One-Way RM ANOVA of alcohol consumption across sessions indicated a significant main effect of session [F _session_ (9, 54) = 34.49, P<0.0001]. Poshoc Dunnet’s test revealed the amount of alcohol consumed during sessions 6-10 when the mice were on LD5001 was significantly higher in comparison to session 1 when the mice were fed TL20I9S. One-way RM ANOVA of alcohol preference also revealed a significant main effect of session [F _session_ (9, 54) = 7.902, p<0.0001]. Poshoc Dunnet’s test showed that alcohol preference was significantly higher during sessions 6-10 than session 1. We also examined average alcohol consumed (**Fig. 2A** inset) and alcohol preference (**Fig. 2B**, inset) per session during the last 4 sessions for each diet. Paired student’s t-test revealed that alcohol consumption and preference were significantly enhanced when the mice were on LD5001 compared to TL2019S) diet. We also examined water consumption across sessions and found no significant differences in water consumption as mice were switched from TL2019S to LD5001 diet (**Fig. S1A**).

**Figure 2:**
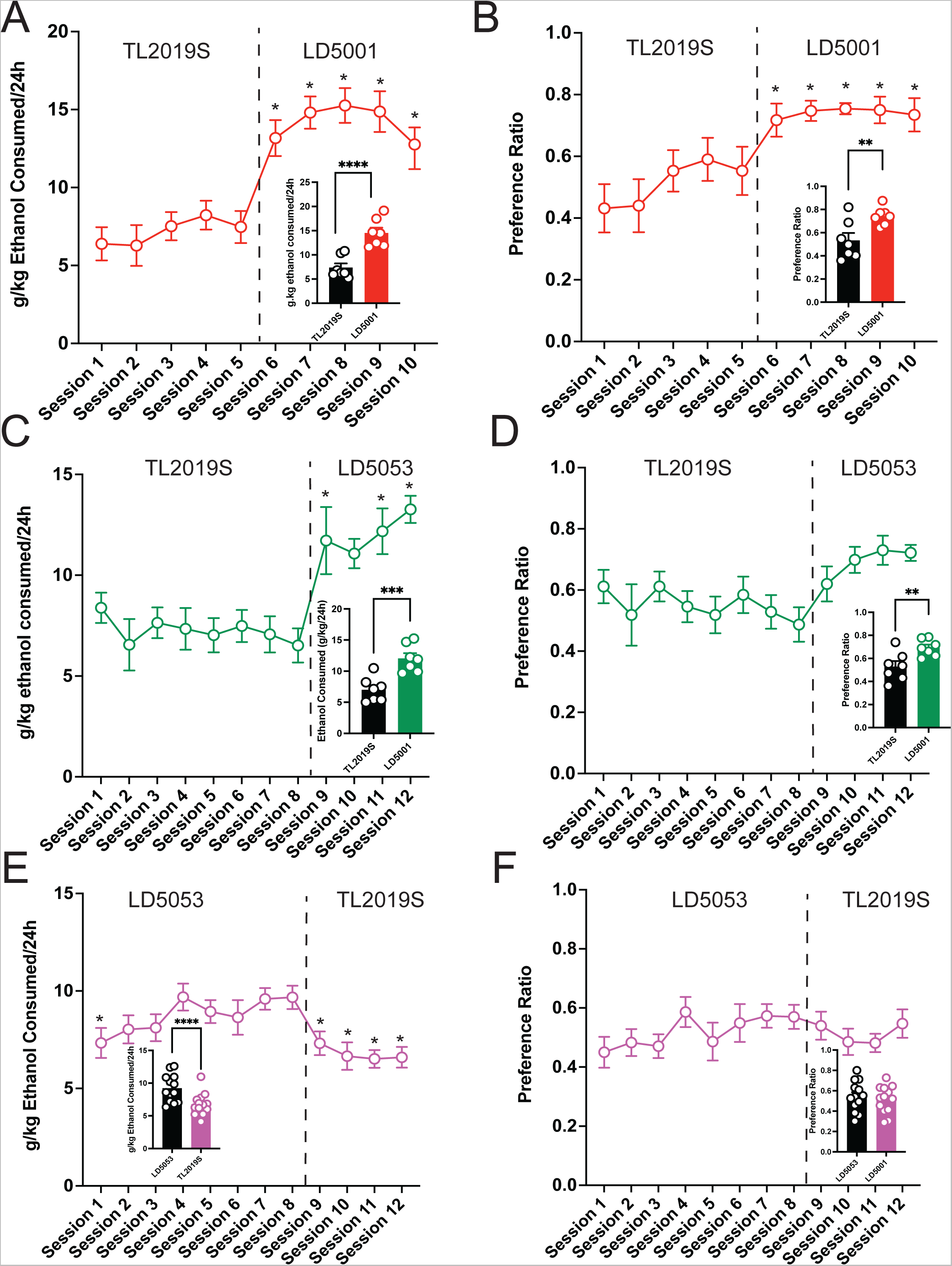
The effects of switching diets on alcohol consumption in males is shown. **A)** Mice consumed significantly more alcohol and showed enhanced preference for alcohol **(B)** when switched from TL2019S to LD5001. *, P<0.05, Dunnett’s post-test compared to Session 1. LD5053 consumed more alcohol (**2A inset**) and showed increased alcohol preference (**2B Inset**) compared to mice on TL2019S during the last 4 sessions of each diet. Student’s t-test, ****, P<0.0001, **, P<0.01, N =7/group. Alcohol consumption (**2C**) and preference (**2D**) was also significantly increased when mice were switched from TL2019S to LD5053. *, P<0.05, Dunnett’s post-test compared to Session 1. Analysis of last 4 days of each diet revealed a significant increase in alcohol consumption (**2C, inset**) and preference (**2D, inset**) when mice were fed LD5053. Student’s t-test, ***, P<0.001, ** P<0.01, N = 7 mice/group. Alcohol consumption and preference was significantly decreased (**Fig 2E and F**) when the diet was switched from LD5053 to TJ2019S. *, P<0.05, Dunnett’s post-test compared to Session 4. Comparison of alcohol consumption and preference during the last 4 drinking sessions for each diet revealed a significant decrease in alcohol consumption (**2E**, **inset**) but no change in preference (**2F, inset**). Student’s t-test, *, P<0.05, N =7/group.

Similarly, we found that switching diets from TL2019S to LD5053 also significant increased alcohol consumption (**Fig. 2C**) and preference (**Fig. 2D**). One-Way RM ANOVA revealed a significant main effect of session [F _session_ (11, 66) = 8.834, P<0.0001]. Posthoc Dunnett’s test indicated that mice consumed significantly more alcohol during sessions 9, 11, and 12 when they were on LD5053 compared to session 1 when they were on TL2019S. Similarly One-way RM ANOVA analysis of alcohol preference revealed a significant main effect of session [F _session_ (11, 66) = 2.860, P = 0.004]. However, posthoc Dunnett’s test failed to reveal any significant differences between alcohol preference in session 1 compared to other sessions. We also examined alcohol consumption and preference during the last 4 sessions for each diet. Paired student’s t-test revealed increased alcohol consumption (**Fig. 2C inset**) and preference (**Fig. 2D, inset**) when mice were fed LD5053 compared to when they were fed TL2019S. Water consumption was not significantly affected when diet was switched from TL2019S to LD5053 (**Fig. S1B**).

We next asked if mice that were drinking high amounts of alcohol on LD5053 would continue to maintain high levels of drinking when their diets were switched to TL2019S. We found that switching diets from LD5053 to TL2019S led to a decrease in alcohol consumption (**Fig. 2E**) and preference (**Fig. 2F**). One-Way RM ANOVA analysis revealed a main effect of alcohol drinking session [F _session_ (11, 132) = 5.601, P < 0.0001]. Posthoc Dunnett’s test revealed that mice consumed significantly lower amounts of alcohol on sessions 1, 9, 10, 11, and 12 compared to session 7 when drinking levels had stabilized. One-Way RM ANOVA analysis of alcohol preference revealed a marginally significant main effect of drinking session [F _session_ (11, 132) = 1.722, P = 0.0749]. We also examined average alcohol consumption and preference per session during the last 4 sessions for each diet. We found a significant decrease in alcohol consumption when mice were switched from LD5053 to TL2019S (**Fig. 2E, inset**). However, we did not see a significant reduction in alcohol preference during the last 4 sessions (**Fig. 2F, insert**).

We next examined if these diet influences on alcohol intake could be extended to females. We found that switching diets from TL2019S to LD5001 significantly increased alcohol consumption (**Fig. 3A**) and preference (**Fig. 3B**) in female mice. One-way RM ANOVA indicated a significant main effect of drinking session [F _session_ (11, 77) = 32.2, P = 0.0001]. Posthoc Dunnett’s test revealed that mice consumed significantly more alcohol on sessions 8-12 when they were on LD5001 compared to session 1 when they were on TL2019S diet. One-Way RM ANOVA of alcohol preference also revealed a significant main effect of drinking session [F _session_ (11, 77) = 20.37, P = 0.0001]. Posthoc Dunnett’s tests indicated that female mice displayed enhanced preference for alcohol on sessions 8-12 compared to session 1. We also compared average alcohol consumed and alcohol preference during the last four sessions for each diet. We found that mice consumed (**Fig. 3A, inset**) significantly more alcohol and showed significantly increased alcohol preference (**Fig. 3B, inset**) when they were on LD5001. We also measured water consumption across sessions and found that in contrast to male mice, female mice consumed significantly less water when switched from TL2019S to LD5001 (**Fig. S1C**). One-way RM ANOVA revealed a main effect of session [F _session_ (11, 77) = 11.20, P < 0.0001]. Posthoc Dunnett’s test revealed that water consumption was significantly lower for sessions 11 and 12 compared to session 1.

**Figure 3:**
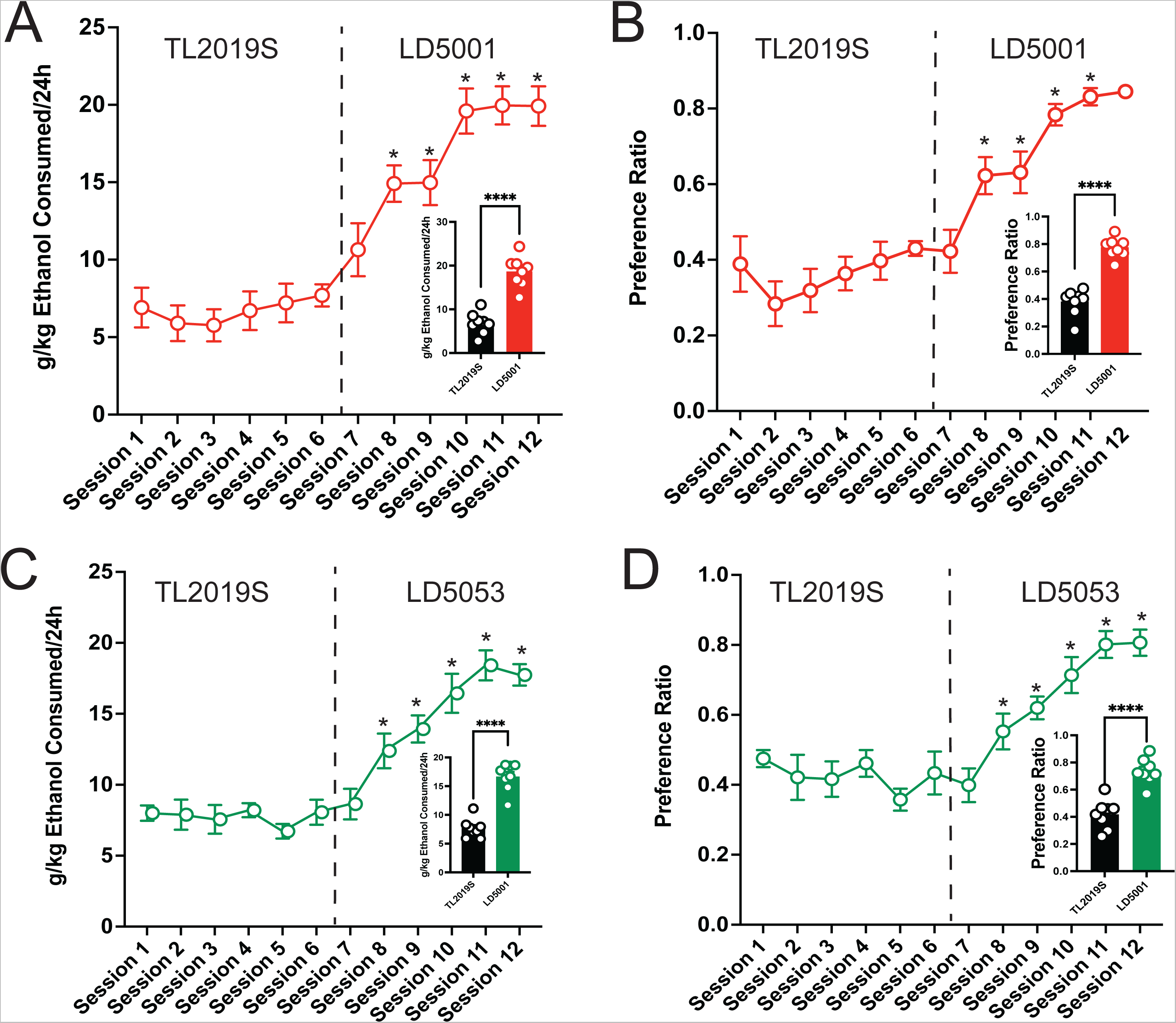
The effects of switching diets on alcohol consumption in females is shown. Female C57BL/6J mice consumed significantly more alcohol **(A)** and showed increased preference **(B)** for alcohol when diet was switched from TL2019S to LD5001. Analysis of alcohol consumption and preference during the last four days on each diet also revealed that female mice consumed more alcohol (**3A, inset**) and showed increased preference (**3B, inset**) for alcohol when fed LD5001. Student’s t-test, ****, P<0.0001, N =8/group. Female mice also consumed significantly (**3C**) more and showed enhanced preference (**3D**) for alcohol when fed LD5053 than when fed TL2019S. Analysis of alcohol consumption during the last 4 sessions for each diet revealed that mice consumed significantly more alcohol (**3C, inset**) and showed more preference for alcohol (**3D, inset**) when fed LD5053 compared to mice fed TL2019S. Student’s t-test, ****, P<0.0001, N = 8/group.

Similarly, female mice also consumed significantly more alcohol when switched from TL2019S to LD5053 (**Fig. 3C**) and preference (**Fig. 3D**). One-Way RM ANOVA revealed a significant main effect of drinking session [F _session_ (11, 77) = 32.56, P = 0.0001]. Posthoc Dunnett’s test revealed that females consumed significantly more alcohol during sessions 8-12 when they were on LD5053 compared to session 1 when they were on TL2019S diet. One-way RM ANOVA of alcohol preference also indicated a significant main effect of drinking session [F _session_ (11, 77) = 20.09, P = 0.0001]. Posthoc Dunnett’s test revealed that mice drank significantly more alcohol during sessions 8-12 when compared to session 1. Examining average alcohol consumed during the last 4 sessions for each diet revealed that female mice on LD5053 consumed significantly more alcohol (**Fig. 3C, inset**) and showed significantly enhanced preference for alcohol (**Fig. 3D, inset**) during the last 4 sessions. Water consumption was also significantly reduced when diet was switched from TL2019S to LD5053 (**Fig. S1D**). One-way RM ANOVA revealed a main effect of session [F _session_ (11, 77) = 9.776, P < 0.0001]. Posthoc Dunnett’s test revealed that water consumption was significantly lower for sessions7, 11 and 12 compared to session 1.

We also examined body weights when the mice were on different diets. Average bodyweights did not differ significantly between males maintained on TL2019S or LD5001 (**Fig. S1A**). There was small but statistically significant increase in bodyweight in mice maintained on LD5053 compared to those maintained on TL2019S (**Fig. S1B**, Student’s-test, P<0.001). There were no significant differences in bodyweights between females maintained on TL2019S and LD5001 (**Fig. S1C**) or LD 5053 (**Fig. S1D**).

We next examined blood ethanol concentrations from male mice that were maintained on the three different diets. Tail vein blood was collected 3h after alcohol bottles were introduced. One-Way ANOVA revealed a significant main effect of diet on alcohol consumption (**Fig. 4**) [F _diet_ (2, 30) = 5.723, P = 0.0078]. Dunnett’s posthoc test indicated that mice on LD5053 and LD5001 showed significantly higher BEC’s than mice on TL2019S which is consistent with these mice consuming higher amounts of alcohol.

**Figure 4:**
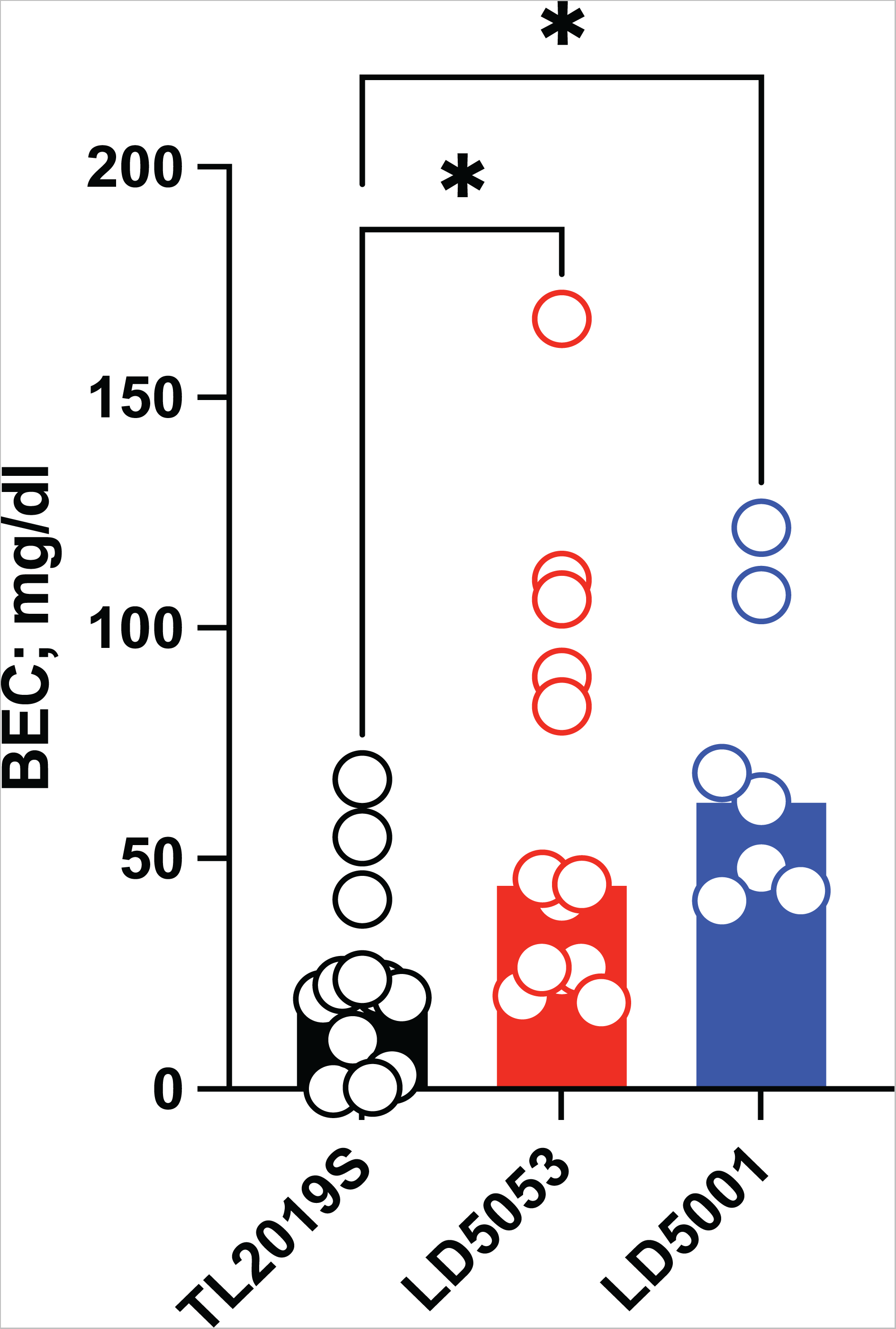
Significantly higher BECs were detected in mice fed LD 5001 and LD5053-fed mice compared to TL2019S. *. = P<0.05, Dunnett’s test, N = 7-13/group.

We saw the highest amount of alcohol consumption in mice that were fed LD5001. Hence, we focused on this diet for future experiments. We next determined if the increased alcohol consumption observed with LD5001 was due to altered taste perception. Therefore, we examined sucrose (4%), saccharin (0.04 and 0.06%), and quinine consumption (100, 175, and 200 μM) in mice maintained on LD5001 or TL2019S diets. We first examined sucrose consumption followed by consumption of increasing concentrations of saccharin and quinine. We found no significant differences in sucrose, quinine, or saccharin consumption between mice that were on TL2019S compared to those that were on LD5001 (**Fig. 5**) suggesting that the increase in alcohol consumption observed in mice that were fed LD5001 is unlikely due to altered taste perception. We next determined if alcohol consumption in LD500-fed mice was sensitive to disruption by quinine adulteration. We added increasing concentrations of quinine to the alcohol. Quinine adulteration dose-dependently decreased alcohol consumption in both TL2019S and LD 5001-fed mice (**Fig. 6**). However, at the highest most aversive concentration of quinine tested, the magnitude of decrease in drinking was greater in TL2019S-fed mice than in mice fed LD5001 indicating that LD5001 led to a higher degree of quinine resistance in these mice (**Fig. 6**). Two-Way RM ANOVA revealed a significant main effect of quinine concentration [F _quinine_ (3, 36) = 24.02, P<0.0001] and a quinine X diet interaction [F _quinine x diet_ (3, 36) = 2.902, P = 0.0481]. Posthoc Sidak test indicate that mice that were fed TL2019S diet consumed significantly less alcohol when it was adulterated with 500μM quinine than mice on LD5001 diet.

**Figure 5:**
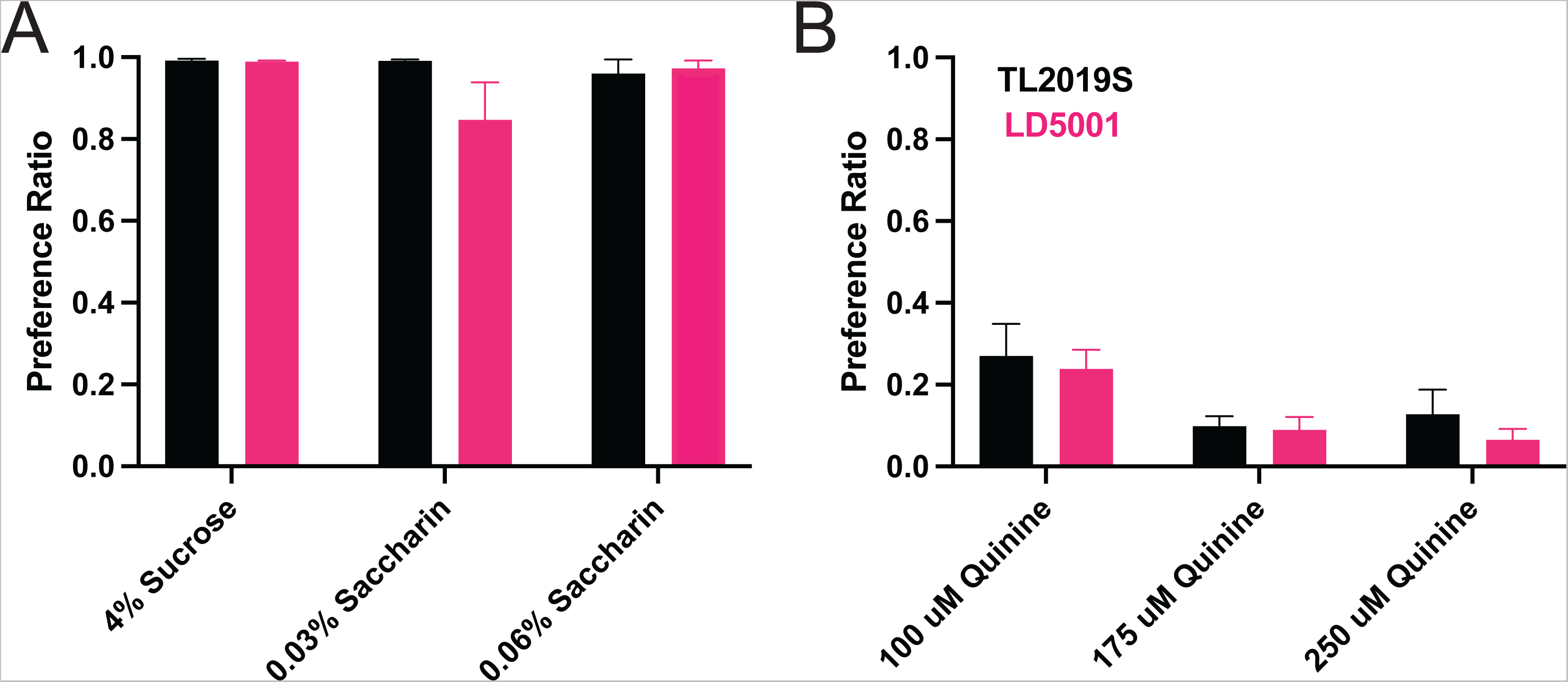
Taste perception was unaltered in mice consuming LD5001. Sucrose, saccharin preference **(A)** and quinine preference **(B)** was unaltered in mice fed with LD5001 compared to TL2019S. N=9-11/group.

**Figure 6:**
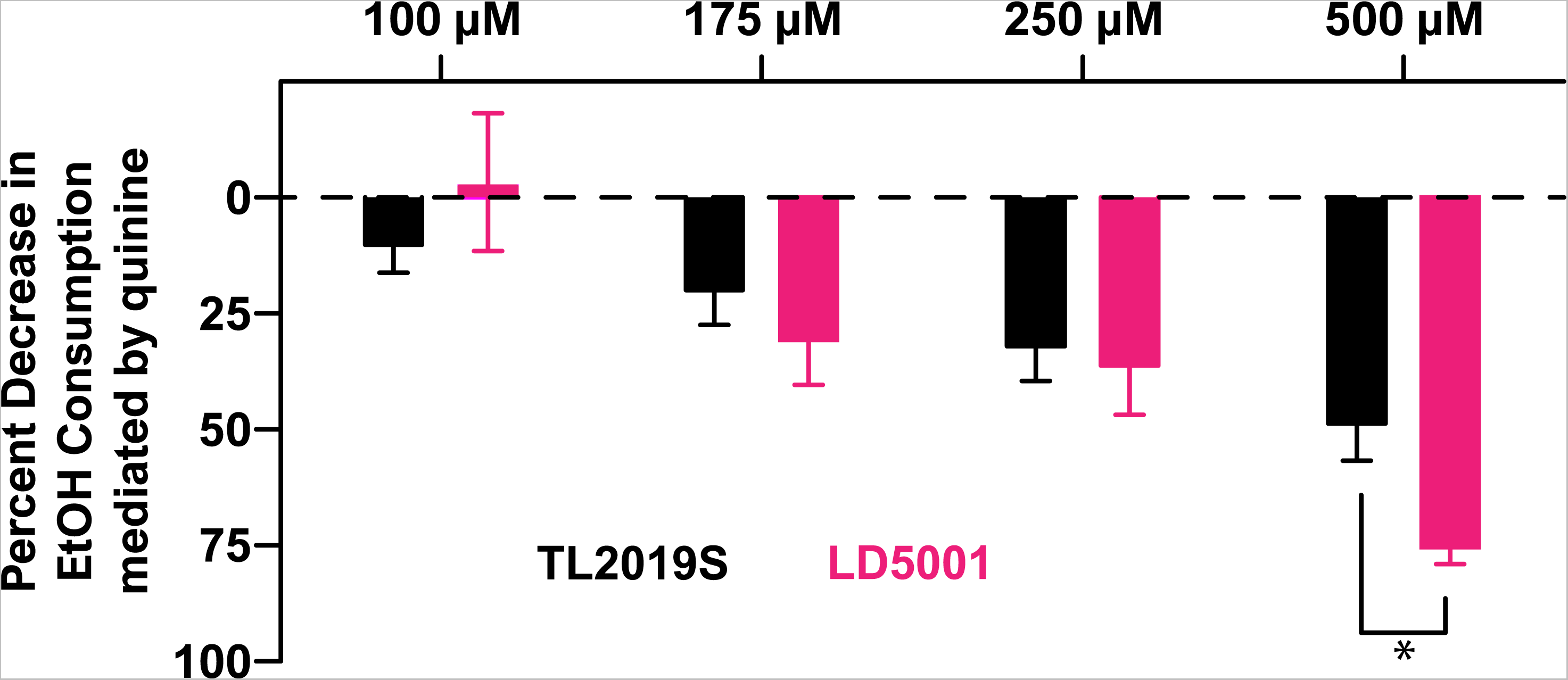
Diet-induced increases in alcohol consumption is resistant to quinine adulteration. Quinine adulteration reduced alcohol consumption in both LD5001 and TL2019S fed mice. However, at the highest most aversive dose of quinine (500 μM), LD5001 fed mice displayed more resistance to quinine adulteration and maintained high levels of intake. *, P<0.05, Sidak post-test. N =7/group.

We next measured the effects of diet on gut microbiome alpha-diversity. Alpha-diversity measures the ecological diversity within-sample, and it can be sub-divided into evenness and richness. Evenness or relative abundance is a measure of the sequence variants uniformity, while sample richness represents the number of each operational taxonomic units (OTUs) present in the sample. We assessed alpha diversity of the stool samples from mice on TL2019S, LD5053, and LD5001 diets. Stool samples were collected from alcohol drinking mice that were part of the diet-switching experiment shown in **Fig. 2**. Mice were switched from TL2019S to LD5053 or LD5001. Samples were collected at two time points: 1) immediately before the last session of alcohol drinking on TL2019S diet and 2) immediately before the 4^th^ alcohol drinking session after the diet switch when mice are on LD5001 or LD5053 diet. Samples were also collected from a separate cohort of water drinking mice that were exposed to the diets for a similar timepoints. **Fig. 7A** shows the effect of diet on relative abundance at the genus level in the water-drinking group. Two-Way ANOVA of relative abundance in water drinking animals revealed a significant diet X genera interaction [F_diet x genera_ (38, 360) = 18.67, P = 7.84E-64]. Posthoc Tukey test showed that mice on TL2019S diet had significantly higher abundance of *Dubosiella* (6.9%) compared to those on LD5053 (0.1%) and LD5001 (0%) (P= 0.0041). Animals on LD5053 showed increased relative abundance of the genus *Alistipes* (31.6%) compared to TL2019S (14.7%) and LD5001 (0.2%) (P<0.0001, Tukey test). Whereas animals on LD5001 had higher relative abundance of the genus *Lachnospiraceae* NK4A136 group (41.4%) than TL2019S (28.4%) and LD5053 (25.8%) (P=0.0047, Tukey test) and higher relative abundance of Bacteroides (24.5%) compared to TL2019S (16.6%) and to LD5053 (18.5%) (P=0.0349, Tukey test). All the relative abundance comparisons for the water-drinking group are compiled in **Table S1.**

**Figure 7:**
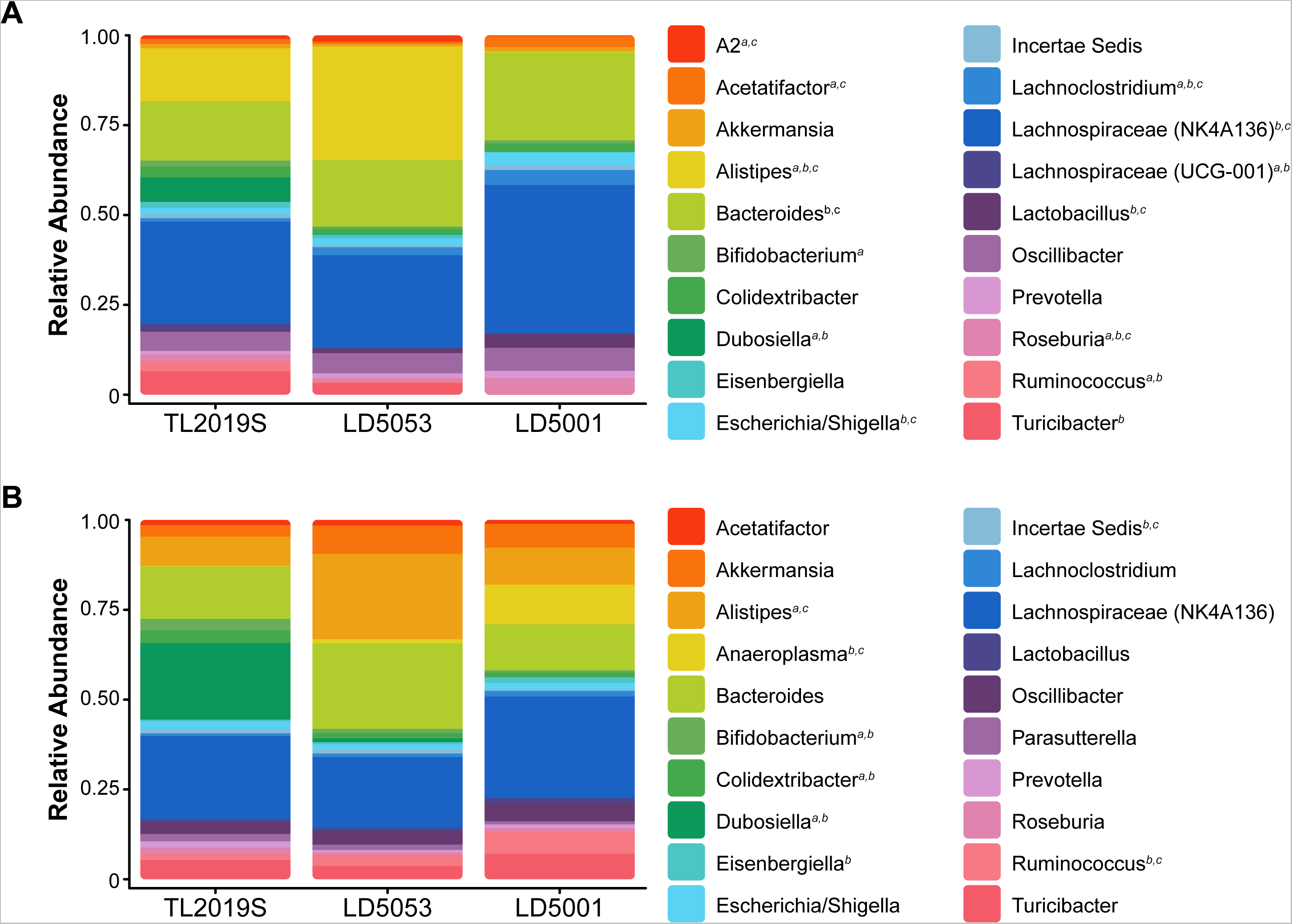
Rodent diets shape gut bacterial abundance. The relative abundances were estimated for the three diets during the water-drinking period **(A)** and after the alcohol sessions **(B)**. N= 7/group.

We next assessed bacterial relative abundance in samples from animals after IA alcohol consumption (**Fig. 7B**). Two-Way ANOVA analysis of relative abundance in alcohol drinking animals revealed a significant diet X genera interaction [F _diet x genera_ (38, 360) = 8.91, P = 5.62E-33]. Alcohol drinking mice on TL2019S diet showed higher relative abundance of *Dubosiella* (21.4%) compared to LD5053 (1%) and LD5001 (0%) (P<0.0001, Tukey test). Animals on LD5053 diet had increased relative abundance of *Alistipes* (23.8%) in comparison to TL2019S (8.1%) and LD5001 (10.3%) (P=0.0008, Tukey test). LD5001 have higher relative abundance of the genus *Anaeroplasma* (11%) compared to TL2019S (0.02%) and to LD5053 (1.1%) (P=0.0008, Tukey test). All the relative abundance comparisons for the water-drinking group are compiled in **Table S2**.

Sample richness was significantly different between the three different diets in the water drinking group (**Fig. 8A**). One-Way ANOVA revealed significant differences in Observed Features [F (2,18) = 4,22, P = 0.0313]. However, Tukey’s post-test did not show statistical differences between the groups; TL2019S vs LD5053 (P = 0.9975), TL2019S vs LD5051 (P = 0.0502) and LD5053 vs LD5051 (P= 0.0573). No differences were found on Shannon Index [F (2,18) = 2. 463, P = 0.1134]. Simpson index was significantly different between the groups [F (2,18) = 4.877, P = 0.0203]. Tukey’s multiple comparison shown significant differences among the groups TL2019S vs LD5001 (mean difference P= 0.0261 95% CI = [0.005722 to 0.09656]). No significant differences were found between TL2019S and LD5053 (P= 0.0558) nor between LD5053 and LD5001 (P=0.9249). Bacterial richness was evaluated in samples from the three diets groups in the alcohol drinking animals (**Fig.8B**). One-Way ANOVA shows the three diet groups have distinct Observed Features [F (2,18) = 3,691, P=0,0454]. No statistical differences were found on the multiple comparison test, TL2019S vs LD5053 (P = 0.0977), TL2019S vs LD5051 (P = 0.9591) and LD5053 vs LD5051 (P = 0.0578). Both Shannon and Simpson Index did not show significant statistical difference between the three diets after alcohol intake.

**Figure 8:**
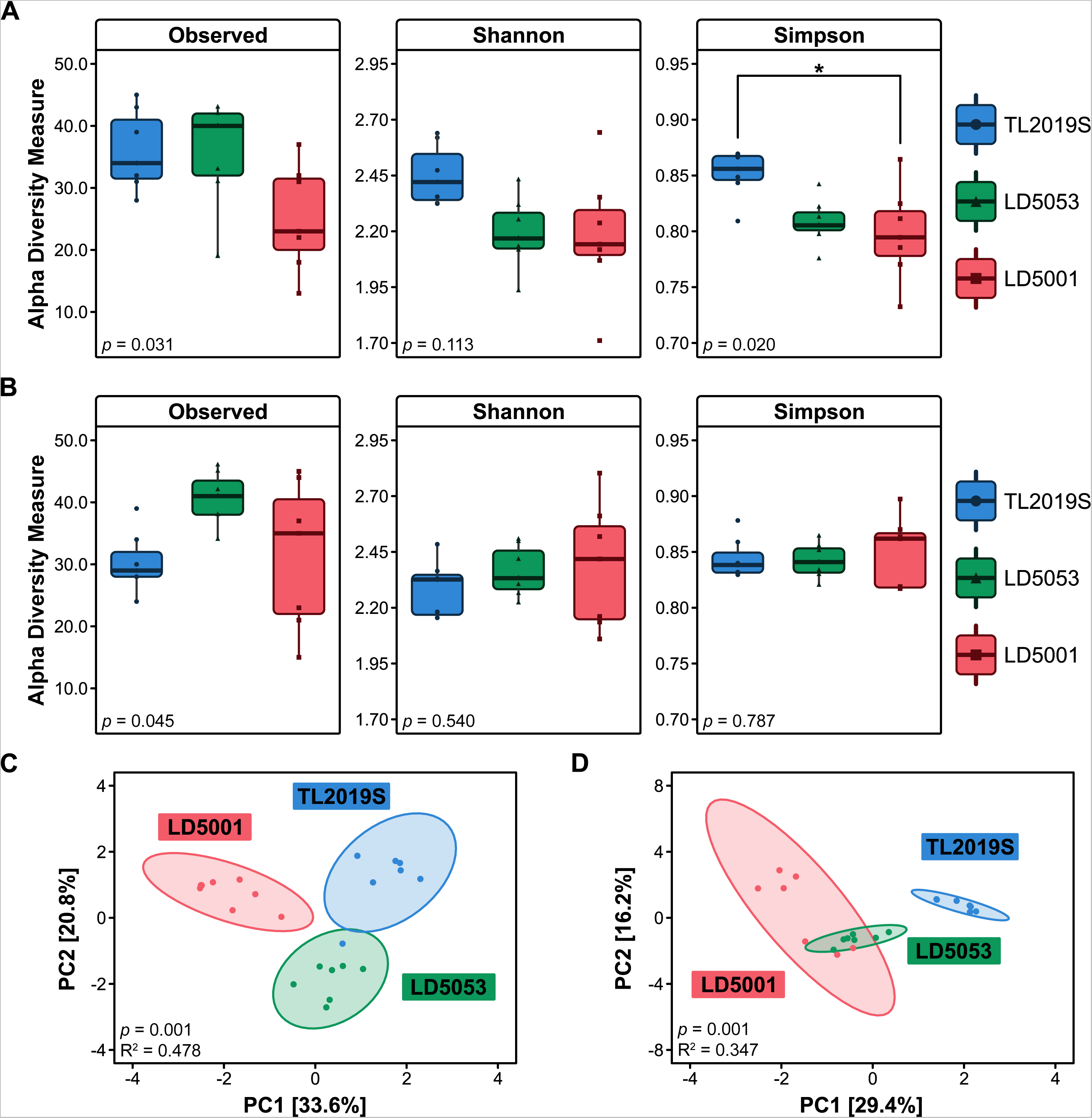
Gut microbial structure is influenced by different chow diets. Bacterial counts were transformed to centered log ratio (CLR) and used to estimate alpha and beta-diversity (N = 7/group). Alpha-diversity was assed using Observed features, Shannon and Simpson indices. **(A)** During the water-drinking sessions, ANOVA test revealed significative differences on Observed features (P=0.031) and Simpson Index (P = 0.020), post-hoc Tukey test only identified significance on Simpson Index between Teklad and LD5001 diets (P = 0.026). **(B)** After intermittent access alcohol (IA) protocol, we did not find significant differences in alpha-diversity. Beta diversity was calculated in terms of Euclidean distance using the centered log ratio (CLR) values. PERMANOVA (ADONIS) results were depicted here as a principal component (PC) graph. **(C)** Beta diversity was significantly changed In the water-drinking group (PERMANOVA [P = 0.001, R2 = 0.478]); *post-hoc* pairwise ADONIS tests revealed significant differences between TL2019s vs LD5053 (P = 0.009); TL2019S vs LD5001 (P = 0.006) and LD5053 vs LD5001 (P = 0.006). **(D)** Animals in IA group also presented significant changes in beta diversity PERMANOVA [P = 0.001, R2 = 0.347]); post-hoc pairwise ADONIS tests revealed significant differences between TL2019S vs LD5053 (P = 0.006); TL2019S vs LD5051 (P = 0.003) and LD5053 vs LD5051 (P = 0.006).

We sought to investigate if the three different diets would result in greater dissimilarities between the bacterial communities, using beta-diversity metrics. Beta diversity was calculated using Aitchison distance, which is the Euclidean distance between data transformed by the centered log-ratio (clr) method. Water drinking mice presented diet-dependent differences in the variation of the communities’ composition. The variance explained by Component 1 was 33.6% and by Component 2 was 20.8% (**Fig. 8C**). ADONIS test revealed a statistically significant difference between the three diets [F (2,18) = 8,24, P = 0.001]. *Post-hoc* pairwise ADONIS tests revealed significant differences between TL2019s vs LD5053 (P = 0.009); TL2019S vs LD5001 (P = 0.006) and LD5053 vs LD5001 (P = 0.006). The beta-diversity values were also significantly different between the alcohol animal groups [F (2,18) = 4,77, P = 0.001] (**Fig. 8D**). Post-hoc pairwise ADONIS tests revealed significant differences between TL2019S vs LD5053 (P = 0.006); TL2019S vs LD5051 (P = 0.003) and LD5053 vs LD5051 (P = 0.006).

## Discussion

Our results indicate that mice fed LD5001 and LD5053 consume more and show enhanced preference for alcohol and attain higher BECs compared to mice fed TL2019S. Sucrose, saccharin, or quinine consumption was not affected by different brands of chow suggesting that the increase in alcohol consumption observed with LD5001 and LD5053 was not due to altered taste perception. Further, alcohol consumption in mice fed LD5001 was more resistant to quinine adulteration when compared with mice that were fed TL2019S. Gut microbiome analysis revealed that the gut bacteria in water and alcohol drinking mice fed LD5001, LD5053, and TL2019S significantly differed in terms of alpha and beta diversity and displayed distinct gut microbial structure.

Previous studies have examined the impact of several commercially available rodent diet formulations on alcohol consumption. One recent study examined binge alcohol consumption in rodents fed LD5001 or TL2920S and found that alcohol consumption and BECs were markedly higher in mice maintained on LD5001 compared to those on TL2920S (Maphis et al., 2022). Mice on LD5001 also displayed increased front-loading behavior and consumed twice as much alcohol in the first 15 minutes than TL2920S-fed mice (Maphis et al., 2022). Another recent study compared mice maintained on four different commercially available rodent diets, LD5001, H7012, H2918, and LDV575 on IA alcohol consumption. They found that mice maintained on LD5001 and H7012 diets consumed high amounts of alcohol compared to the other two diets (Quadir et al., 2020). A previous study looked at the effects of six commercially available rodent diets on alcohol consumption in the “Drinking in the dark” model of binge alcohol consumption and continuous access two bottle-choice drinking. The diets evaluated were RMH3000 (Purina) and Teklad diets T. 2916, T.2918. T.2920x, T.7912, and T.8940. Mice maintained on T.7912 consumed the highest amount and showed the largest preference for alcohol (Marshall et al., 2015). However, none of these studies have examined mechanisms by which these different diets influence alcohol intake.

Consistent with these previously published findings, we found that standard rodent diet formulations can profoundly and differentially influence alcohol consumption in mice. The three diets we tested, LD5001, LD5053, and TL2019S do not differ significantly in macronutrient composition but have numerous differences in micronutrient composition (**Table 1**). The most notable differences were in the amounts of vitamins and minerals with higher amounts of Vitamins A, E, K3, B1, B2 in TL2019S compared to LD5001 and LD5053. Higher amounts of Calcium, Phosphorous, Potassium, Chloride, and Magnesium were present in TL2019S compared to LD5001 and LD5053. There were also higher amounts of Sulfur, Cobalt, Fluorine, and Chromium in LD5001 and LD5053 compared to TL2019S. There were also textural differences between LD5053 and LD5001 compared to TL2019S with TL2019S being grittier in texture than the LD diets. The difference in texture might result from a variety of reasons including the method employed to sterilize these foods. TL2019S is sterilized by autoclaving whereas LD5001 and LD5053 are gamma irradiated. It is unclear which of these differences contribute significantly to increased alcohol consumption.

One recent metanalysis examined the concentration of isoflavones in various commercially available rodent diet formulations and found that they may vary as much as 20-600mg/g of diet. There is a significant positive correlation between alcohol consumption and preference and isoflavone concentration in mice (Eduardo and Abrahao, 2022). We do not know the isoflavone concentrations in the rodent diets that were used in our study and hence it remains to be determined if isoflavones underlie the differences observed in our study.

Although we didn’t directly measure the amount of food consumed in our study, we found no differences in bodyweights when the mice were maintained on different diets, which suggests that perhaps food intake was not significantly different. We also found that water intake was significantly decreased in male mice maintained on LD5053 and in females maintained on both LD5001 and LD5053 compared to mice maintained on TL2019S. One possible reason for this could be that water consumption was reduced to compensate for the increased amounts of alcohol consumed.

We did not find any diet-induced changes in sucrose, saccharin, or quinine preference in our study. This finding contrasts with a previous study suggesting diet formulations that increase alcohol consumption also increase sucrose/saccharin consumption (Marshall et al., 2015). An interesting finding in our study is that the diet that resulted in the highest amount of alcohol consumption, LD5001, also resulted in significant resistance to quinine adulteration of alcohol compared to mice on TL2019S diet. Quinine-resistant alcohol intake is frequently used to model drinking despite aversive consequences (De Oliveira Sergio et al., 2023). It is important to point out that both TL2019S- and LD5001-fed mice decreased their drinking when alcohol solution was adulterated with increasing concentrations of quinine. However, the magnitude of this reduction was greater in TL2019S-fed mice than in LD5001-fed mice at the highest concentration of quinine tested, suggesting increased resistance to quinine adulteration in LD5001-fed mice. These results suggest that rodent diet formulation can influence the development of compulsive alcohol seeking in mice.

To determine the mechanism by which diet could influence alcohol intake, we examined gut microbiota in water- and alcohol-drinking mice on different diets. Diet can profoundly influence gut microbial composition (Tuck et al., 2020), which can in turn influences alcohol consumption and reward via multiple mechanism including signaling through the gut-brain axis (Leclercq et al., 2019; Leclercq et al., 2020; Quoilin et al., 2023; Salavrakos et al., 2021). In fact, a recent study showed that fecal microbiota transplant from healthy donors can reduce alcohol preference and craving in people with AUD, and this behavior is transmissible to germ-free mice (Wolstenholme et al., 2022).

Animals fed with TL2019S had lower alcohol preference and consumption and higher abundance of the genus *Dubosiella* compared to the two other diets. Interestingly, *Dubosiella* was previously associated with the production of beneficial metabolites, such as short chain fatty-acids (SCFA) (Kadyan et al., 2023). SCFAs such as acetate, butyrate and propionate supplementation is effective in reducing stress-induced gut-brain axis disorders (van de Wouw et al., 2018). The higher abundance of *Dubosiella* and its metabolites might be associated with lower motivation to alcohol drinking. The genus *Alistipes* was markedly increased in water-drinking mice that were fed LD5053, and this characteristic was maintained after alcohol exposure. Prior research demonstrated high abundance of this genus in mice subjected to stress (Bangsgaard Bendtsen et al., 2012) and in patients diagnosed with depression (Naseribafrouei et al., 2014). *Alistipes* hydrolyze tryptophan to produce indole. Tryptophan is also precursor for serotonin, so higher abundance of *Alistipes* could indirectly reduce serotonin availability in the gut, impairing gut-brain axis signaling (Vlainić et al., 2016) which could in turn influence alcohol consumption and preference.

The gut microbiome of LD5001-fed mice showed the greatest changes in relative abundance after alcohol consumption. In water drinking mice maintained on LD5001, *Lachnospiraceae* NK4A136 and *Bacteroides* were the predominant genera. The relative abundance of these genera was significantly higher in LD5001-fed mice compared to TL2019S- and LD5053-fed mice. Increased abundance of *Lachnospiraceae* NK4A136 is associated with elevated stress, anxiety, and depression-like behaviors (Pizarro et al., 2021). Additionally, the abundance of this bacteria predicts lower concentrations of serotonin in the prefrontal cortex (PFC) (McGaughey et al., 2019; Wong et al., 2016). It is possible that the increased alcohol consumption we observe in mice fed LD5001 could be due to increased anxiety-like behaviors, although we did not test for anxiety-like behaviors in this study. Bacteroides is a genus of commensal microbes, and it is usually shown to be depleted after alcohol exposure (Addolorato et al., 2020; Xiao et al., 2018); however, the association of Bacteroides and alcohol-seeking behavior in not known and needs to be further investigated.

One caveat with our study is that unlike alcohol-drinking mice, the water-drinking mice were maintained on TL2019S, LD5053 or LD5001 and not subject to diet switching. Hence, it is possible that some of the changes in bacterial diversity observed in alcohol-drinking mice after the diet switch may not be observed in mice that were maintained on one type of diet throughout the experiment.

Gut microbiota could modulate alcohol consumption in a variety of ways including by altering neuroinflammation, myelin synthesis, blood brain barrier permeability, and production of metabolites that alter signaling along the gut-brain axis (Leclercq et al., 2014; Leclercq et al., 2019). In this study we did not perform metabolomics studies to determine if the different diets altered the metabolic profile of the gut microbiota. The goal of this study was limited to identifying bacterial genera that could account for diet-induced differences in alcohol consummatory behaviors. Hence one limitation of this study is that we did not establish causal relationships between specific bacterial genera and alcohol consumption. Future studies will employ techniques such as fetal microbial transplant in germ-free mice and metabolomics to determine specific bacterial genera and bacterial metabolite that mediate changes in alcohol intake.

In summary, our results provide strong evidence that standard rodent diet formulations can profoundly influence alcohol consumption and preference, as well as the development of compulsive like alcohol consumption. Hence, it is imperative that studies examining voluntary alcohol consumption document the type of standard rodent diet that the mice were maintained on to increase reproducibility across labs. Importantly, our result also suggests that commercially available rodent diet formulations can profoundly and differentially impact gut microbiome diversity which could contribute to regulating alcohol consummatory behaviors.

## Acknowledgements

This work was funded by NIH grant 1R01AA0227293 and LSUHSC startup finds to RM. SW was supported by a medical student alcohol research internship (T35AA021097-10). The authors would like to thank Robert Siggins and for many helpful comments.

## Bibliography

Addolorato, G. et al. (2020) Gut microbiota compositional and functional fingerprint in patients with alcohol use disorder and alcohol-associated liver disease. Liver Int, 40, 878–888.

Avegno, E.M. & Gilpin, N.W. (2019) Inducing Alcohol Dependence in Rats Using Chronic Intermittent Exposure to Alcohol Vapor. Bio Protoc, 9, e3222.

Bangsgaard Bendtsen, K.M., et al. (2012) Gut microbiota composition is correlated to grid floor induced stress and behavior in the BALB/c mouse. PLoS One, 7, e46231.

Bravo, J.A. et al. (2011) Ingestion of Lactobacillus strain regulates emotional behavior and central GABA receptor expression in a mouse via the vagus nerve. Proc Natl Acad Sci U S A, 108, 16050–16055.

By, I.M.P.A.C.T.T.I. (2022) Beta-diversity distance matrices for microbiome sample size and power calculations - How to obtain good estimates. Comput Struct Biotechnol J, 20, 2259–2267.

Chang, C.S. & Kao, C.Y. (2019) Current understanding of the gut microbiota shaping mechanisms. J Biomed Sci, 26, 59.

Crabbe, J.C. & Wahlsten, D. (2003) Of mice and their environments. Science, 299, 1313–1314.

De Oliveira Sergio, T., Frasier, R.M. & Hopf, F.W. (2023) Animal models of compulsion alcohol drinking: Why we love quinine-resistant intake and what we learned from it. Front Psychiatry, 14, 1116901.

Dubinkina, V.B. et al. (2017) Links of gut microbiota composition with alcohol dependence syndrome and alcoholic liver disease. Microbiome, 5, 141.

Eduardo, P.M.C. & Abrahao, K.P. (2022) Food composition can influence how much alcohol your animal model drinks: A mini-review about the role of isoflavones. Alcohol Clin Exp Res, 46, 6–12.

García-Cabrerizo, R. et al. (2021) Microbiota-gut-brain axis as a regulator of reward processes. J Neurochem, 157, 1495–1524.

Glantz, M.D. et al. (2020) The epidemiology of alcohol use disorders cross-nationally: Findings from the World Mental Health Surveys. Addict Behav, 102, 106128.

Hwa, L.S. et al. (2011) Persistent escalation of alcohol drinking in C57BL/6J mice with intermittent access to 20% ethanol. Alcohol Clin Exp Res, 35, 1938–1947.

Kadyan, S. et al. (2023) Resistant starches from dietary pulses modulate the gut metabolome in association with microbiome in a humanized murine model of ageing. Sci Rep, 13, 10566.

Koob, G.F. & Volkow, N.D. (2016) Neurobiology of addiction: a neurocircuitry analysis. Lancet Psychiatry, 3, 760–773.

Leclercq, S. et al. (2014) Role of inflammatory pathways, blood mononuclear cells, and gut-derived bacterial products in alcohol dependence. Biol Psychiatry, 76, 725–733.

Leclercq, S. et al. (2020) Gut Microbiota-Induced Changes in β-Hydroxybutyrate Metabolism Are Linked to Altered Sociability and Depression in Alcohol Use Disorder. Cell Rep, 33, 108238.

Leclercq, S. et al. (2019) The gut microbiota: A new target in the management of alcohol dependence. Alcohol, 74, 105–111.

Lohoff, F.W. (2022) Targeting Unmet Clinical Needs in the Treatment of Alcohol Use Disorder. Front Psychiatry, 13, 767506.

Lynch, S.V. & Pedersen, O. (2016) The Human Intestinal Microbiome in Health and Disease. N Engl J Med, 375, 2369–2379.

Maiya, R. et al. (2021) Differential regulation of alcohol consumption and reward by the transcriptional cofactor LMO4. Mol Psychiatry, 26, 2175–2186.

Maphis, N.M., Huffman, R.T. & Linsenbardt, D.N. (2022) The development, but not expression, of alcohol front-loading in C57BL/6J mice maintained on LabDiet 5001 is abolished by maintenance on Teklad 2920x rodent diet. Alcohol Clin Exp Res, 46, 1321–1330.

Marshall, S.A. et al. (2015) Assessment of the Effects of 6 Standard Rodent Diets on Binge-Like and Voluntary Ethanol Consumption in Male C57BL/6J Mice. Alcohol Clin Exp Res, 39, 1406–1416.

McGaughey, K.D. et al. (2019) Relative abundance of Akkermansia spp. and other bacterial phylotypes correlates with anxiety- and depressive-like behavior following social defeat in mice. Sci Rep, 9, 3281.

Nagendra, H. (2002) Opposite trends in response for the Shannon and Simpson indices of landscape diversity. Applied geography, 22, 175–186.

Naseribafrouei, A. et al. (2014) Correlation between the human fecal microbiota and depression. Neurogastroenterol Motil, 26, 1155–1162.

Pizarro, N. et al. (2021) Sex-Specific Effects of Synbiotic Exposure in Mice on Addictive-Like Behavioral Alterations Induced by Chronic Alcohol Intake Are Associated With Changes in Specific Gut Bacterial Taxa and Brain Tryptophan Metabolism. Front Nutr, 8, 750333.

Quadir, S.G. et al. (2020) Effect of different standard rodent diets on ethanol intake and associated allodynia in male mice. Alcohol, 87, 17–23.

Quoilin, C. et al. (2023) Exploring the links between gut microbiota and excitatory and inhibitory brain processes in alcohol use disorder: A TMS study. Neuropharmacology, 225, 109384.

Rhodes, J.S. et al. (2005) Evaluation of a simple model of ethanol drinking to intoxication in C57BL/6J mice. Physiol Behav, 84, 53–63.

Salavrakos, M. et al. (2021) Microbiome and substances of abuse. Prog Neuropsychopharmacol Biol Psychiatry, 105, 110113.

Stärkel, P. et al. (2018) Intestinal dysbiosis and permeability: the yin and yang in alcohol dependence and alcoholic liver disease. Clin Sci (Lond), 132, 199–212.

Tabakoff, B. & Hoffman, P.L. (2000) Animal models in alcohol research. Alcohol Res Health, 24, 77–84.

Tordoff, M.G. et al. (2002) The maintenance diets of C57BL/6J and 129X1/SvJ mice influence their taste solution preferences: implications for large-scale phenotyping projects. J Nutr, 132, 2288–2297.

Tuck, C.J. et al. (2020) Nutritional profile of rodent diets impacts experimental reproducibility in microbiome preclinical research. Sci Rep, 10, 17784.

van de Wouw, M. et al. (2018) Short-chain fatty acids: microbial metabolites that alleviate stress-induced brain-gut axis alterations. J Physiol, 596, 4923–4944.

Vengeliene, V., Bilbao, A. & Spanagel, R. (2014) The alcohol deprivation effect model for studying relapse behavior: a comparison between rats and mice. Alcohol, 48, 313–320.

Vlainić, J.V. et al. (2016) Probiotics as an Adjuvant Therapy in Major Depressive Disorder. Curr Neuropharmacol, 14, 952–958.

Wolstenholme, J.T. et al. (2022) Reduced alcohol preference and intake after fecal transplant in patients with alcohol use disorder is transmissible to germ-free mice. Nat Commun, 13, 6198.

Wong, M.L. et al. (2016) Inflammasome signaling affects anxiety- and depressive-like behavior and gut microbiome composition. Mol Psychiatry, 21, 797–805.

Xiao, H.W. et al. (2018) Gut microbiota modulates alcohol withdrawal-induced anxiety in mice. Toxicol Lett, 287, 23–30.

