## Supplemental data for "Standard rodent diets differentially impact alcohol consumption and preference and gut microbiome diversity"

**Figure S1:** The effects of switching diets on water consumption is shown. There were no significant differences in water consumption when diet was switched from TL2019S to LD 5001 **(A)** or LD5053 **(B)** in males. N =7/group. In females, water consumption was significantly reduced when diet was switched from TL2019S to LD5001. *, P<0.05, Dunnett's post-test compared to Session 1 **(C)**. Water consumption was also significantly reduced when diet was switched from TL2019S to LD5053 diet **(D)**. *, P<0.05, Dunnett's post-test compared to session1. N =8/group.

**Figure S2**: Average bodyweights across sessions during the diet switching experiment is shown. shown. Bodyweights did not significantly differ in male mice when they were maintained on LD5001 when compared to when they were maintained on LD 5053 **(A)**. There was a small but significant increase in body weight in mice maintained on LD5053 compared to when they were maintained on TL2019S **(B)**. ***, P<0.001, paired t-test, N =7/group. Bodyweights were not significantly different in females maintained on LD5001 **(C)** or LD5053 **(D)** when compared to mice maintained on TL2019S. N =8/group.

### Figure S1

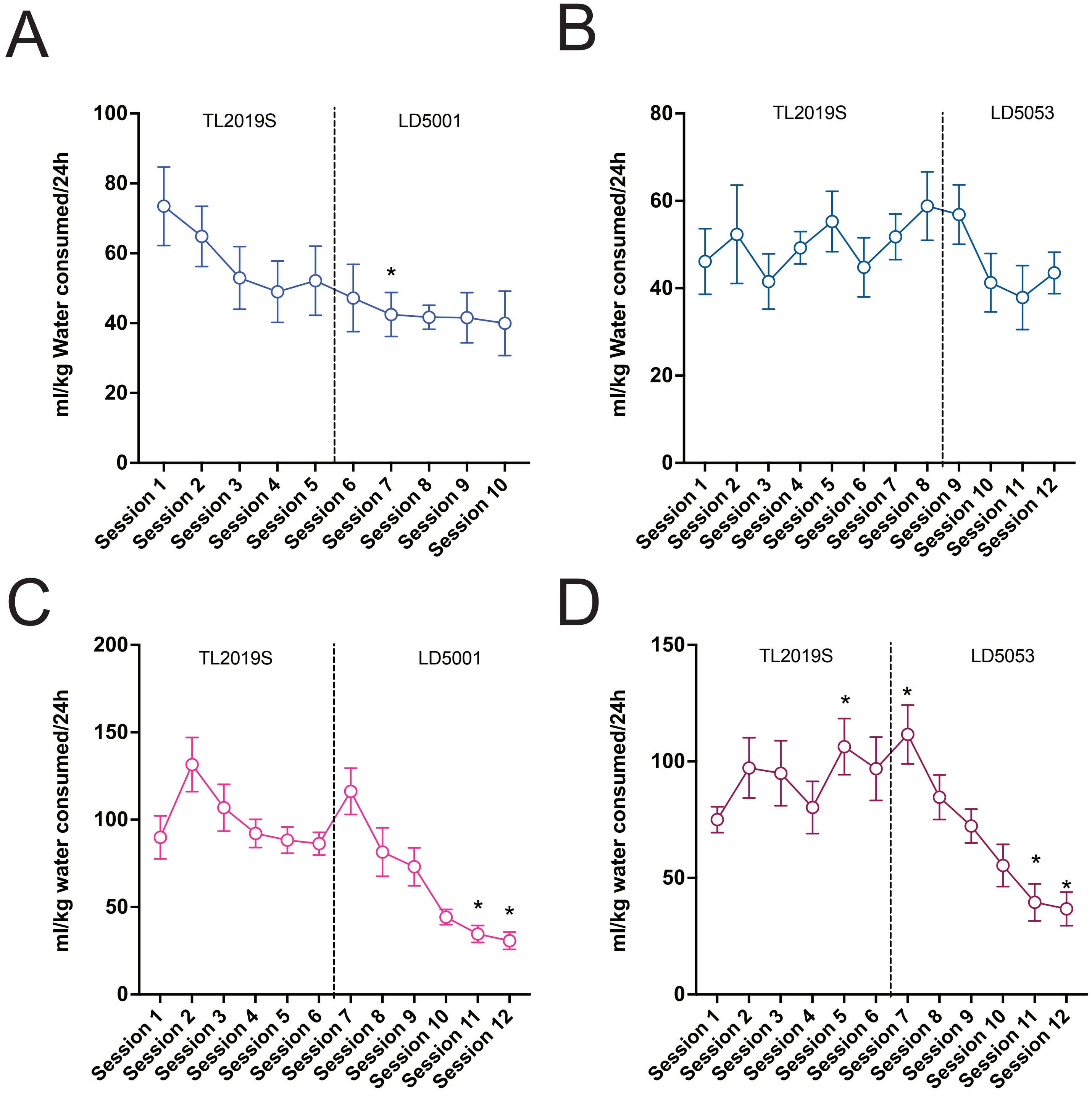
A
B
C
D
TL2019S
LD5001
LD5053
ml/kg Water consumed/24h
ml/kg water consumed/24h
Session 1
Session 2
Session 3
Session 4
Session 5
Session 6
Session 7
Session 8
Session 9
Session 10
Session 11
Session 12
*

Figure S2

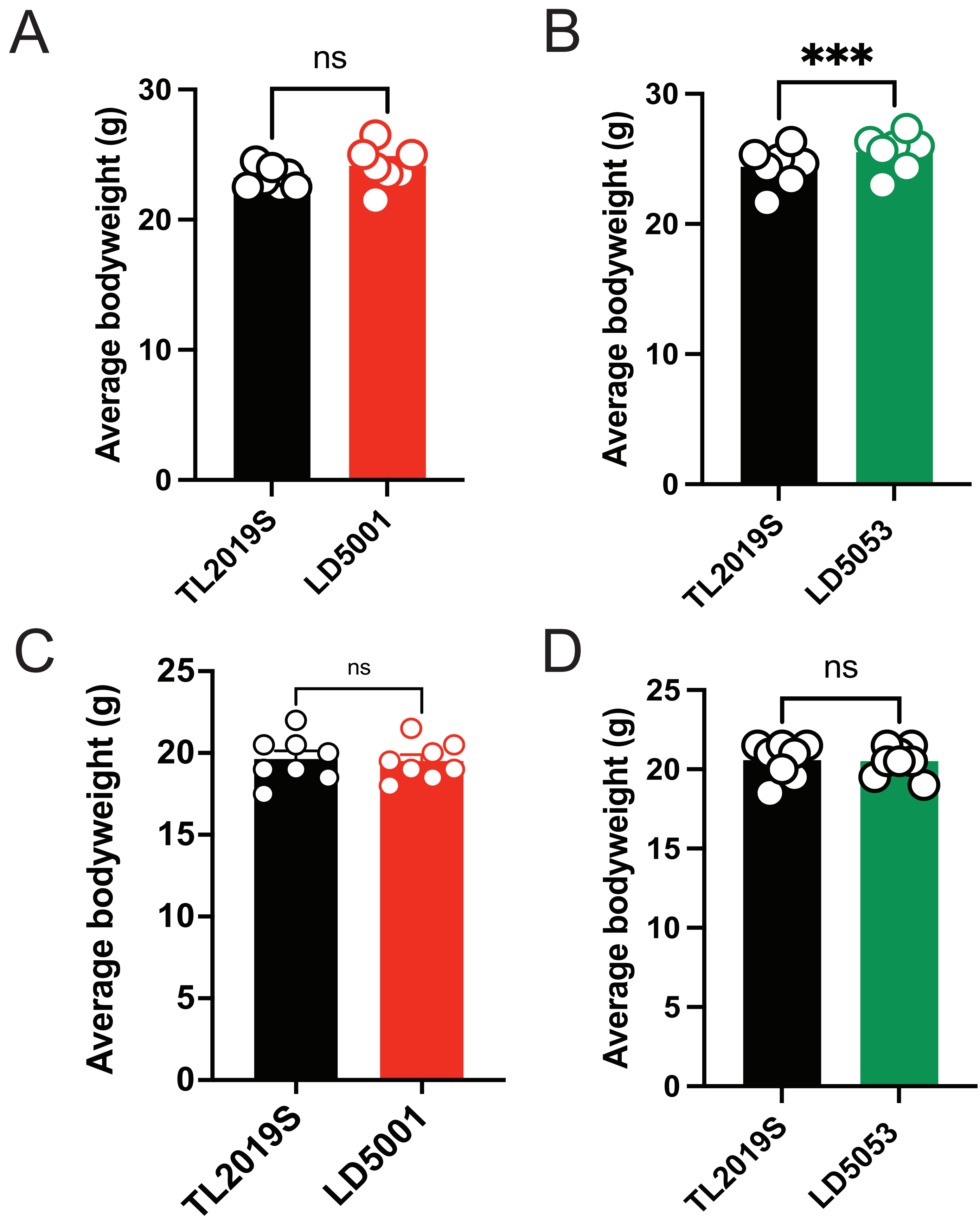
A
ns
Average bodyweight (g)
30
20
10
0
TL2019S
LD5001
B
***
Average bodyweight (g)
30
20
10
0
TL2019S
LD5053
C
ns
Average bodyweight (g)
25
20
15
10
5
0
TL2019S
LD5001
D
ns
Average bodyweight (g)
25
20
15
10
5
0
TL2019S
LD5053

**Table S1: Water Relative Abundance, p values obtained from TukeyHSD tests on significant genera**

| Genus | Comparison | | |
|---|---|---|---|
| | TL2019S-LD5053 | TL2019S-LD5001 | LD5053-LD5001 |
| A2 | 5.00E-02 | 5.66E-01 | 5.70E-03 |
| Acetatifactor | 1.46E-01 | 2.47E-02 | 3.39E-04 |
| Alistipes | 4.17E-07 | 1.14E-06 | 1.23E-11 |
| Bacteroides | 9.84E-01 | 3.43E-02 | 4.85E-02 |
| Bifidobacterium | 1.60E-02 | 7.89E-02 | 7.15E-01 |
| Dubosiella | 3.80E-03 | 3.44E-03 | 9.99E-01 |
| Escherichia/Shigella | 9.97E-01 | 2.26E-02 | 2.64E-02 |
| Lachnoclostridium | 2.58E-02 | 6.63E-06 | 2.76E-03 |
| Lachnospiraceae_NK4A136_group | 1.00E+00 | 4.54E-03 | 4.61E-03 |
| Lachnospiraceae_UCG-001 | 1.33E-02 | 3.66E-03 | 8.25E-01 |
| Lactobacillus | 4.44E-01 | 6.30E-06 | 7.70E-05 |
| Roseburia | 3.36E-02 | 1.59E-04 | 7.30E-07 |
| Ruminococcus | 1.55E-03 | 1.51E-04 | 5.41E-01 |
| Turicibacter | 1.62E-01 | 2.74E-02 | 6.28E-01 |

P value adjusted with Benjamini & Hochberg adjustment

**Table S2: Alcohol Relative Abundance, p values obtained from TukeyHSD tests on significant genera**

| Genus | Comparison | | |
|---|---|---|---|
| | TL2019S-LD5053 | TL2019S-LD5001 | LD5053-LD5001 |
| Alistipes | 2.85E-04 | 8.25E-01 | 1.04E-03 |
| Anaeroplasma | 9.29E-01 | 2.51E-03 | 5.60E-03 |
| Bifidobacterium | 1.65E-03 | 8.96E-05 | 3.86E-01 |
| Colidextribacter | 3.87E-03 | 1.58E-03 | 9.13E-01 |
| Dubosiella | 5.68E-08 | 3.14E-08 | 9.25E-01 |
| Eisenbergiella | 7.15E-01 | 1.49E-02 | 7.41E-02 |
| Incertae_Sedis | 6.49E-01 | 7.39E-03 | 4.84E-02 |
| Ruminococcus | 1.14E-01 | 3.61E-05 | 3.39E-03 |

P value adjusted with Benjamini & Hochberg adjustment
